# A macrophage-based screen identifies antibacterial compounds selective for intracellular *Salmonella* Typhimurium

**DOI:** 10.1101/383828

**Authors:** Michael J. Ellis, Caressa N. Tsai, Jarrod W. Johnson, Shawn French, Wael Elhenawy, Steffen Porwollik, Helene Andrews-Polymenis, Michael McClelland, Jakob Magolan, Brian K. Coombes, Eric D. Brown

## Abstract

*Salmonella* Typhimurium (*S*. Tm) evades the innate immune response by residing within host phagocytes. To identify inhibitors of intracellular *S*. Tm growth, we performed parallel chemical screens against *S*. Tm growing in macrophage-mimicking media and within macrophages. These screens identified novel antibacterials, and revealed that antibiotics with limited Gram-negative coverage are active against intracellular *S*. Tm. Screening of a *S*. Tm deletion library in the presence of one compound, metergoline, revealed that outer membrane perturbation enhanced activity *in vitro*. Combined with our observation of atypical cell surface characteristics of intracellular *S*. Tm, our work indicates that the bacterial outer membrane is permeabilized within macrophages. We show that metergoline targets the bacterial cytoplasmic membrane, and prolongs animal survival during a systemic *S*. Tm infection. This work highlights the predictive nature of intracellular screens for *in vivo* efficacy, and uncovers new aspects of bacterial physiology of intracellular *S*. Tm.

## Introduction

The stagnant antibiotic discovery pipeline is most concerning for intracellular infections. Bacterial pathogens that survive within phagocytic cells manipulate host signaling to resist the antimicrobial activity of the immune system and form persister cells that tolerate antibiotic treatment^1,2^. As such, intracellular infections are often recurrent and persistent, warranting an increase in antimicrobial research specific for such pathogens and an improved understanding of the genetic requirements for intracellular survival. Intracellular bacteria occupy modified phagosomes (*Salmonella, Mycobacterium, Francisella*), inclusions (*Chlamydia*), lysosomes (*Legionella, Coxiella*) or the cytosol (*Listeria, Shigella, Ricksettia*) of host cells, which offer considerably different environments relative to standard nutrient-rich growth media^3^. Emerging evidence suggests that non-canonical intracellular bacteria (*Staphylococcus aureus*, *Streptococcus pneumoniae*) are also able to survive within host cells^4,5^. In these intracellular environments, genes that are otherwise dispensable for growth in nutrient-rich media often become essential, constituting a novel antimicrobial target space that is currently under explored^6^. Genes conditionally essential within host cells may be overlooked in experimental systems that do not resemble the intracellular environment; indeed, recent systematic studies of the genetic requirements for growth in infection-relevant conditions have revealed additional essential genes relative to those required for growth *in vitro*^7–9^. High-throughput screening platforms in conditions that closely resemble the intracellular environment have the potential to uncover novel antimicrobials that target conditionally essential genes.

*Salmonella enterica* serovar Typhimurium (*S*. Tm) is a facultative intracellular pathogen and one of the leading causes of gastroenteritis worldwide^10^. Enteric salmonellosis is typically self-limiting in healthy individuals, although in the elderly and immunocompromised, *S*. Tm invades and manipulates professional phagocytes into reservoirs for systemic spread through the reticuloendothelial system^11^. *Salmonella* infections are commonly treated with fluoroquinolones and macrolides, both of which penetrate mammalian cells efficiently to reach the bacterial target; however, resistance to both of these classes is increasing^12^. Of further concern are extensively drug-resistant *S*. Typhi^13^ and a highly invasive, multidrug-resistant variant of *S*. Tm that first emerged in sub-Saharan Africa (ST313)^14^. The emergence of antibiotic resistance across clades of *Salmonella* species threatens to intensify an already significant global health burden, underscoring the importance of novel antibiotic drug discovery.

During infection, *S*. Tm occupies macrophages and neutrophils within modified phagosomes called Salmonella-containing vacuoles (SCVs). A predominantly intracellular lifestyle affords protection from extracellular host immune pressures, including bile salts, antimicrobial peptides, and serum complement^11^. Host cells also act as reservoirs for dissemination to systemic sites, and often provide unique metabolic environments to shelter intracellular pathogens from nutrient competition with resident bacteria^15^. Several antimicrobial mechanisms intrinsic to the intracellular environment (metal depletion, vacuolar acidification, oxidative stress)^16^ serve as environmental signals for multiple two-component regulatory systems in *S*. Tm that detect and respond to immune stresses with alterations in virulence gene expression^17^.

In line with previous work suggesting discordance between *in vitro* and *in vivo* drug susceptibility^18^, we speculated that *S*. Tm displays altered sensitivity to antimicrobials within macrophages. In particular, genes or processes that become essential within the intracellular environment may represent new drug targets or even sensitize *S*. Tm to existing or novel antibiotics. Accordingly, we screened an *S*. Tm ordered gene deletion collection^19^ for growth impairment in macrophage-mimicking media. These data and subsequent experiments with bacteria internalized in macrophages revealed that *S*. Tm is sensitized to the intracellular environment following genetic perturbation of metabolic and cell envelope biogenesis pathways. Indeed, a high-throughput compound screen against intracellular *S*. Tm in macrophages identified several small molecules with conditional efficacy in only acidic minimal media or macrophages, and not in nutrient-rich media. Interestingly, we also observed potentiation of canonically Gram-positive targeting antibiotics in the intracellular environment, which we ascribe to a loss of normal outer membrane structure in macrophage-internalized *S*. Tm. This screen led to the identification of an intracellular-selective antimicrobial, metergoline, which perturbs the bacterial cytoplasmic membrane and prolongs animal survival in a murine model of systemic *S*. Tm infection.

## Results

In the systemic phase of infection, different populations of *S*. Tm encounter varying degrees of nutrient limitation and immune stressors, whether internalized within macrophages and neutrophils, persisting extracellularly in the bloodstream, or invading intestinal epithelial cells. We sought to identify growth-inhibitory small molecules that are specific for intramacrophage *S*. Tm, as macrophages are one of the primary host cell types manipulated by *Salmonella* spp. for replication and systemic dissemination. We reasoned that to be selective for intracellular bacteria, a compound should interfere with one or more biological processes that are required only for growth in this environment. We therefore wanted to survey the genetic requirements for *S*. Tm growth in cultured macrophages.

### Nutrient biosynthesis and cell envelope maintenance are required for intracellular growth of *S*. Tm

While others have identified *Salmonella* genes that become required (i.e. conditionally essential) for growth in conditions mimicking those *in vivo*^20-24^; there has been no systematic, genome-scale survey of the impact of gene deletion on *S*. Tm survival in cultured macrophages. The *S*. Tm str. 14028S genome contains ~4200 non-essential genes, ~3700 of which have been deleted in the ordered *Salmonella* single-gene deletion (SGD) collection^19,25^. Traditional macrophage infection assays are not amenable to high-throughput screening with this large a number of individual strains, so we first aimed to identify mutants with impaired growth in macrophage-mimicking media. Low-phosphate, low-magnesium media (LPM) is acidic, nutrient-poor, limited in Mg^2+^/PO_4_^3-^ ion availability, and has been established to mimic the SCV and alter gene expression in *S*. Tm^26^. Recent work has suggested that LPM and other types of host-mimicking growth media are highly predictive of antimicrobial efficacy *in* vivo^18^. As LPM has been accepted as a suitable approximation for the intracellular environment within host cells, we measured growth of each strain in the SGD collection in this medium (Figure 1A, Supplementary Dataset S1). Notably, approximately half of 125 SGD mutants impaired for growth in LPM were also impaired for intracellular replication in RAW264.7 macrophages, 25 of which had a ≥30% replication defect compared to WT (Figure 1B, Supplementary Dataset S2). Interestingly, the genes important for growth in LPM and intracellular replication are primarily involved in nutrient biosynthesis and metabolism, as well as biological processes related to cell envelope homeostasis. Among others, this includes amino acid (e.g. *aroE, hisA, argH, argG, serB*) and nucleotide (e.g. *pyrF, pyrE, purG, purE, purF*) biosynthesis, as well as LPS modifications and maintenance (e.g *rfc, pgm, rfaK, rfaH, rbK, rfaI, pgm*). Taken together, these data suggest that compounds that target metabolism or the bacterial cell surface would be effective against intracellular *S*. Tm.

**Figure 1.**
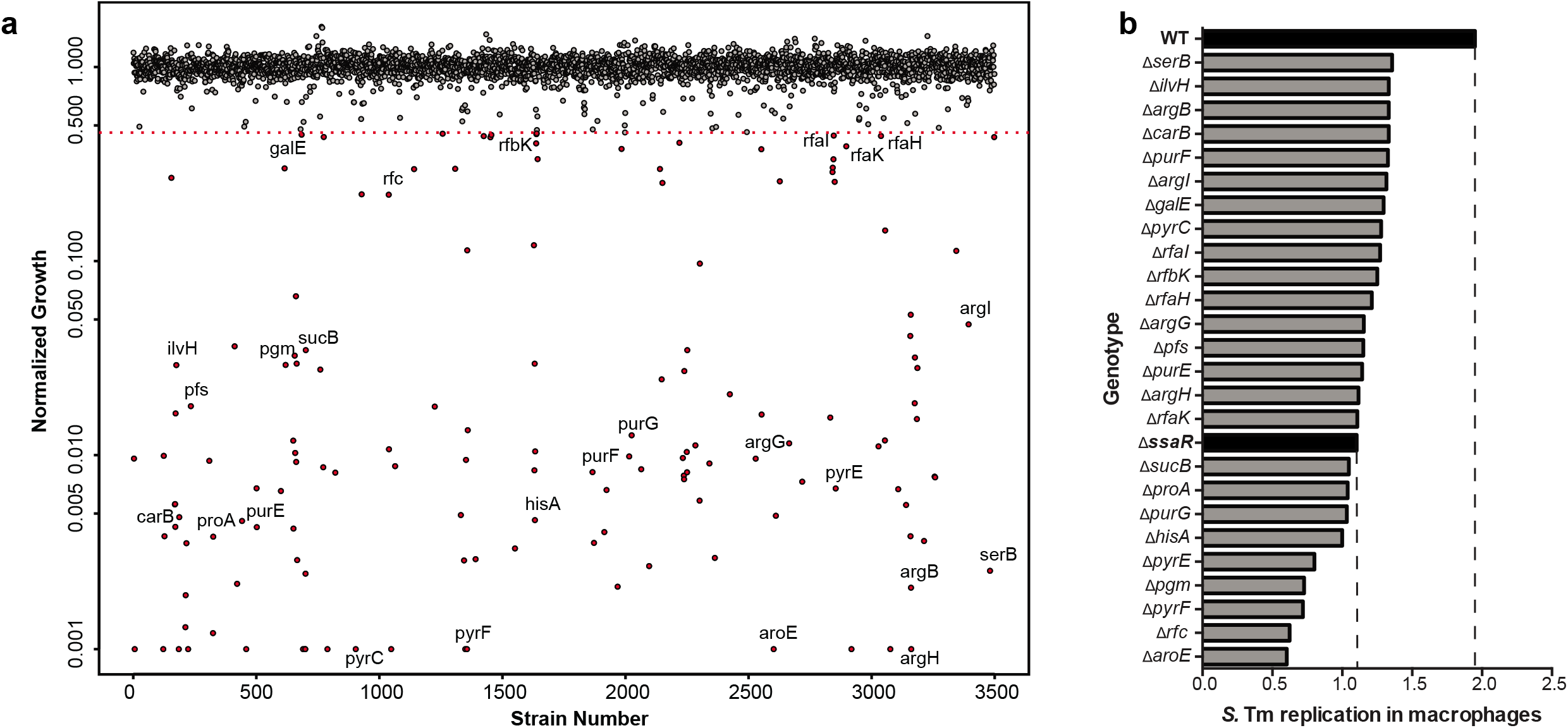
Genetic requirements for *S*. Tm growth in host-mimicking media and macrophages. (A) Index plot showing normalized growth of mutant strains from the *Salmonella* single-gene deletion (SGD) collection in LPM media, sorted in order of chromosomal position of deleted genes. Values shown per strain represent the calculated mean growth of three replicate screens, normalized to account for plate and positional effects. Points below the red dotted line represent genes with growth values less than 3.5σ from the mean of the dataset. Strains that exhibited low growth and were used in follow-up experiments are labeled. (B) Replication of selected mutant strains from the SGD collection in RAW264.7 macrophages over 7 hours. A ∆*ssaR* mutant strain was used as a low replication control, highlighted in black. Replication values reflect the mean of technical duplicates.

### Identification of small molecule inhibitors of intracellular *S*. Tm

To identify putative antimicrobials effective against intracellular *S*. Tm, we conducted two parallel chemical screens with an annotated compound collection composed largely of previously-approved drugs (Supplementary Table 1). We screened *S*. Tm grown in (i), LPM and (ii), RAW264.1 macrophages (Figure 2A). The collection of 1600 chemicals used in these screens includes ~250 known antibacterial compounds with defined targets in Gram-positive and -negative bacteria. To perform the intracellular screen, we modified the standard gentamicin protection assay used to measure bacterial replication in macrophages. Briefly, we measured luminescence from a constitutively expressed luciferase reporter gene to enable *in situ* estimations of intramacrophage bacterial viability (Figure S1). Together, these high-throughput screens identified 130 compounds with growth-inhibitory activity (Supplementary Dataset S3), ~40% of which were effective against both macrophage-internalized bacteria and cells grown in LPM. The overlap in hits between these screens was consistent with ~50% concordance between our genetic screens in LPM and macrophages.

**Figure 2.**
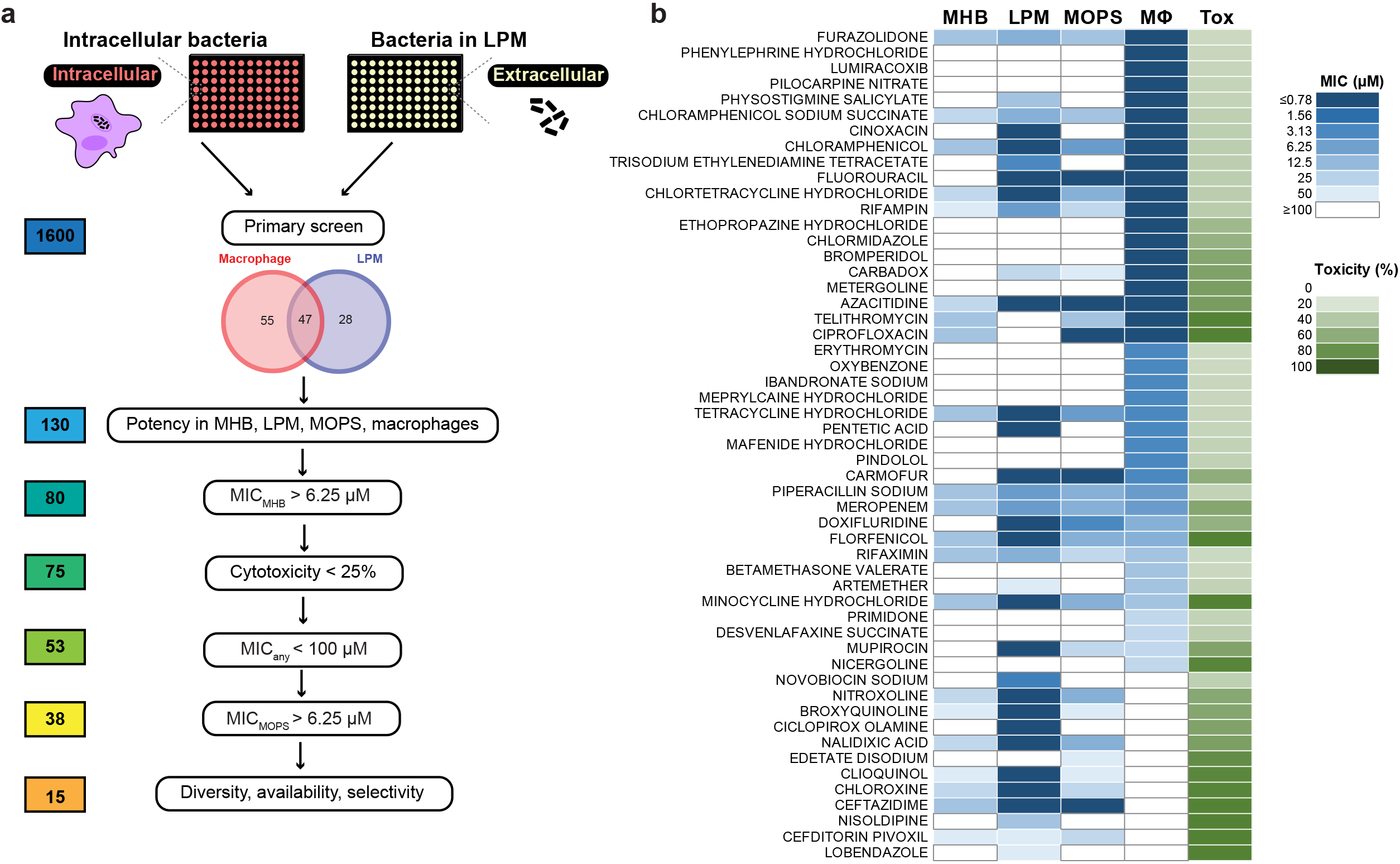
Chemical screen identifies novel compound activities against intracellular *S*. Tm. (A) Screening workflow to identify compounds with intracellular antimicrobial activity against *S.* Tm grown in acidic LPM media and internalized in RAW264.7 macrophages. Secondary screening pipeline is shown below with the number of compounds remaining at each step to the left. (B) Potency and toxicity analysis of all primary screen actives represented as a heat map. Shown are the minimum inhibitory concentrations (MIC) for all compounds against *S*. Tm grown in MHB, LPM, MOPS (OD_600_) and inside RAW264.7 macrophages (MΦ, luminescence); the final column (Tox) reports cytotoxicity of 50 μM compound after 2 hours exposure of RAW264.7 macrophages, determined via lactate dehydrogenase release. Values shown reflect the mean of duplicate measurements.

We next analyzed the potency of all 130 actives from the primary screen. Conditions included *S*. Tm grown in LPM, RAW264.1 macrophages, as well as standard nutrient-rich (cation-adjusted Mueller-Hinton Broth, MHB) and nutrient-poor (MOPS glucose minimal media, MOPS) microbiological media. We also measured lactate dehydrogenase (LDH) release from uninfected macrophages as a measure of toxicity and did not pursue compounds with toxicity >25% (percentage relative to maximum LDH release, see Materials and Methods) (Figure 2B, Supplemental Dataset S4). Using a cutoff for minimum inhibitory concentration (MIC) of 6.25 μM, we eliminated compounds with activity in MHB, as these are least likely to be intramacrophage-selective *in vivo*. We found that nucleoside analogs (e.g. doxifluridine, fluorouracil, azacitidine, carmofur) were effective against intracellular *S*. Tm and bacteria grown in LPM or MOPS minimal media, but not MHB. As genes involved in nucleotide biosynthesis are required for intracellular growth (Figure 1, e.g. *pyrC, pryE, pyrF, purE, purF, purG*), these data suggest that our genetic and chemical screens accurately probed the intramacrophage environment encountered by *S*. Tm. Our analysis identified 31 compounds that were exclusively active in LPM or macrophages and not in MHB or MOPS. We prioritized 15 of these, based on low host cell cytotoxicity, potency, chemical diversity and commercial availability.

Our work thus far utilized a luminescence-based readout for bacterial viability and an immortalized macrophage cell line. We tested the 15 priority actives for the ability to reduce intracellular replication (measured by CFU enumeration) of *S*. Tm in primary bone marrow-derived macrophages isolated from C57BL/6 mice (Figure 3A). Ciprofloxacin is an antibiotic routinely used in salmonellosis infection treatment and was therefore included for comparison.

**Figure 3.**
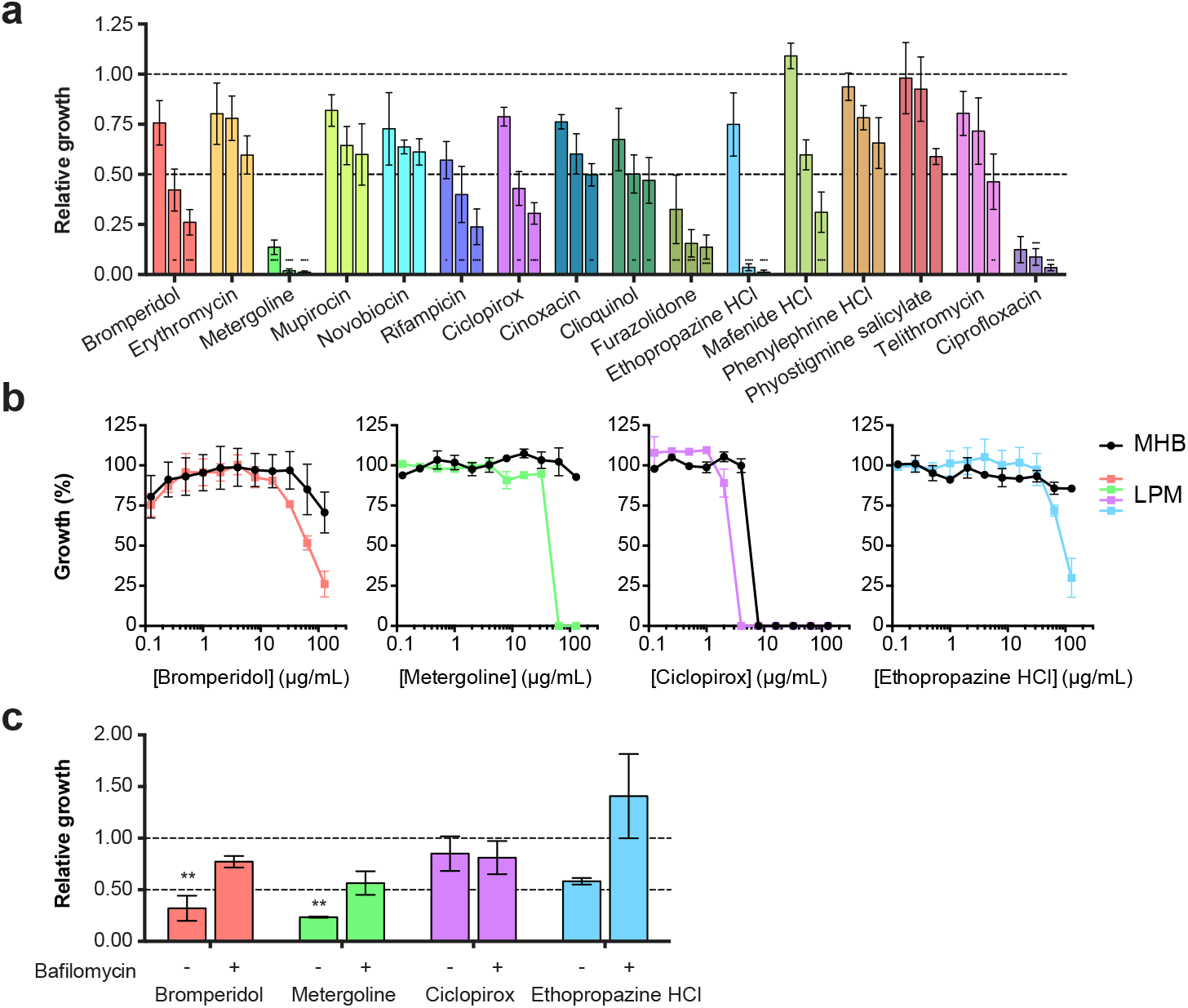
Confirmation of growth inhibitory activity of intracellular-active compounds. (A) Intracellular *S*. Tm replication measured in primary bone marrow-derived macrophages (BMMs) isolated from C57BL/6 mice. Relative growth reflects replication over 4 hours, normalized to bacterial growth in BMMs treated with DMSO. Compounds were added at 8, 64, and 128 μg/mL, as shown in increasing concentrations for each. Bars show the mean and SEM of three independent biological replicates. (B) Potency analysis of *S*. Tm growth inhibition for bromperidol, metergoline, ciclopirox, ethopropazine. Growth is normalized to a DMSO control (set to 100%). (C) Intracellular *S*. Tm replication measured in primary BMMs isolated from C57BL/6 mice, pre-treated with either bafilomycin (10 μM) or DMSO. Relative growth reflects replication over 4 hours, normalized to bacterial growth in BMMs treated with DMSO. Compounds were added at 8 μg/mL. Values in (A) and (C) were compared via two-way ANOVA with Bonferroni correction for multiple testing. *P<0.05, **P<0.01, ***P<0.001, ****P<0.0001.

Of the compounds tested, 11 significantly reduced bacterial replication within 4 hours at a concentration of 128 μg/mL (P <0.05, two-way ANOVA, Bonferroni multiple test correction). We prioritized those which reduced bacterial viability >2-fold at a concentration of 64 μg/mL or less: bromperidol, metergoline, rifampicin, ciclopirox, furazolidone, and ethopropazine. Rifampicin and furazolidone are known antibiotics and therefore were not investigated further. The four remaining compounds had increased activity in LPM relative to MHB (Figure 3B), suggesting selectivity of these antimicrobials for an *in vivo*-mimicking environment. We measured the impact of these four compounds on bacterial replication in the presence of bafilomycin, an inhibitor of vacuolar ATPase activity that blocks acidification of the SCV. Remarkably, bafilomycin pretreatment of bone marrow-derived macrophages antagonized the activity of bromperidol, metergoline, and ethopropazine administered at 8 μg/mL (Figure 3C), suggesting that their antimicrobial efficacy was dependent on the acidic intracellular environment. Taken together, our data suggest that these compounds work in concert with one or more components of the intracellular environment. Metergoline was the most potent of these compounds and was the focus of subsequent studies of mechanism and *in vivo* efficacy.

### The Gram-negative outer membrane antagonizes metergoline activity

Metergoline significantly reduced *S*. Tm growth within primary macrophages at 8 μg/mL. The MIC of metergoline against *S*. Tm grown in MHB is above its solubility limit (~256 μg/mL), although its MIC in LPM is 128 μg/mL. The discordance in metergoline potency against bacteria grown in MHB, LPM, or within macrophages led us to hypothesize that intracellular bacteria were hypersensitized to metergoline as a result of a unique mechanism-of-action (MOA). To gain insight into the antibacterial activity of metergoline, we first used the SGD collection to investigate its genetic interactions. We grew this collection of *S*. Tm deletion mutants in the presence of a sub-lethal concentration of metergoline (100 μg/mL) and compared the growth of each mutant to an MHB control (Figure 4A and Supplementary Dataset S5). These data revealed that deletions of genes involved in assembly of LPS (*rfaG, rfaQ*), outer membrane (OM) integrity (*asmA, tolQ, tolR, yfgL*), turnover of cell wall (*ldcA, prc, nlpI*), or RND efflux pumps (*tolC*, *acrB*) are required for normal growth in the presence of metergoline. We observed that deletions of *tolC, tolR*, and *rfaQ* in *S*. Tm all resulted in hypersensitivity to metergoline in the intracellular environment (Figure S2A). From this, we inferred that: (i) disruption of OM or cell wall integrity increases metergoline potency, and (ii) metergoline is a substrate of the AcrAB-TolC efflux pump. Indeed, RfaG and RfaQ are both required for synthesis of the inner core of LPS, Prc and NlpI are both components of a periplasmic protease complex important for peptidoglycan recycling, TolQ and TolR are components of the Tol-Pal complex required for OM integrity, and AcrB and TolC are inner- and outer-membrane components of the major multi-drug efflux pump in *Enterobactericeae*.

**Figure 4.**
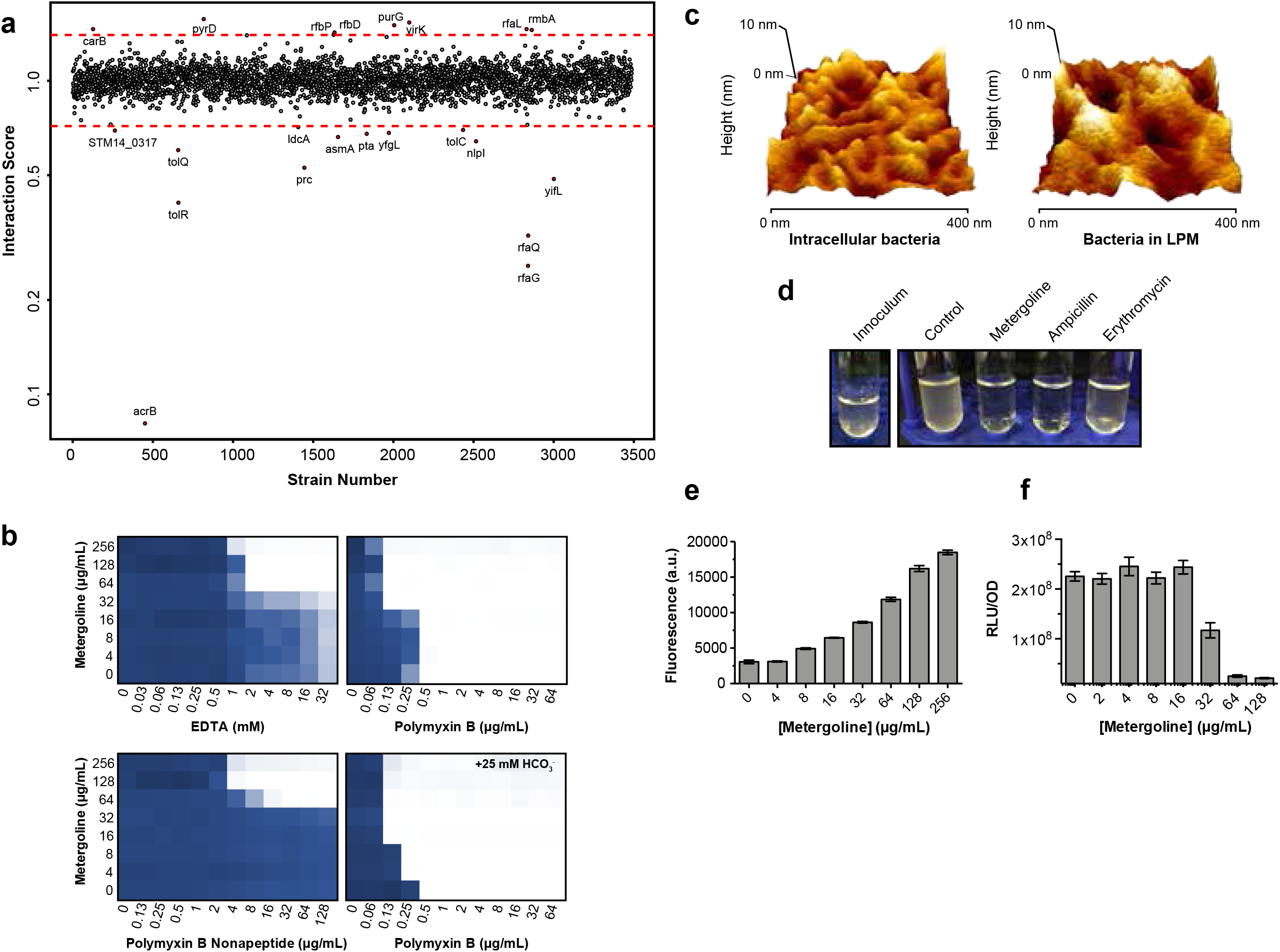
Metergoline targets the inner membrane of outer membrane-perturbed *S*. Tm. (A) Index plot showing sensitivity of SGD collection mutant strains grown in MHB with 100 μg/mL metergoline. Strains are sorted based on chromosomal position of the deleted gene. The chemical-genetic interaction score was calculated by dividing normalized growth of each mutant in the presence of metergoline divided by normalized growth in MHB. Red lines indicate 3σ from the mean of the dataset and values represent the mean from duplicate screens. (B) Chequerboard broth microdilution assay showing dose-dependent potentiation of metergoline by membrane-perturbing agents against *S*. Tm grown in MHB. Where indicated, sodium bicarbonate was added to media at a final concentration of 25 mM. Higher growth is indicated in dark blue and no detectable growth in white. Results are representative of at least two independent experiments. (C) Atomic force microscopy (AFM) of *S*. Tm isolated from BMMs (left) and grown in LPM (right). Image depicts a representative high-resolution surface scan of a region on a bacterial cell from each condition. Surface topology of the region was recorded and rendered in 3D, illustrating a ~10 nm range within peaks and valleys of the scan. (D) Turbidity of cultures of *S*. Tm ∆*tolC* in MHB after growth to mid-log phase (left, inoculum) then 2.5 hours of growth at 31°C in the presence of metergoline (200 μg/mL), ampicillin (16 μg/mL), erythromycin (16 μg/mL), or a DMSO control. Note that erythromycin is bacteriostatic and culture turbidity did not change relative to the inoculum; ampicillin (bactericidal) and metergoline both cleared culture turbidity. (E) DiSC_3_(5) assay on late-log phase *S*. Tm grown in MHB supplemented with 10 mM EDTA to enable DiSC_3_(5) binding to the inner membrane. Cells were loaded with DiSC_3_(5) prior to a 1 min incubation with increasing concentrations of metergoline; bars show mean of fluorescence values and standard error for two biological replicates. (F) *S*. Tm grown in MHB with 1 mM EDTA to early-log phase, then exposed to metergoline for 30 min. Cellular ATP levels were estimated by luciferase activity (relative light units, RLU) normalized to optical density (OD_600_). Bars show the mean and standard error for two biological replicates.

Given the hypersensitivity of OM mutants to metergoline, we reasoned that OM-perturbing agents might potentiate its activity. In line with this, we noted that in our chemical screens, several antibiotics conventionally active against only Gram-positive bacteria (rifampin, rifamixin, mupirocin, novobiocin) inhibited growth of *S*. Tm in LPM and/or macrophages. These antibiotics are known to be potentiated against Gram-negative bacteria in the presence of OM perturbing agents (e.g. colistin^27^ and pentamidine^28^), leading us to speculate that acidic (LPM) or intracellular (macrophage) environments compromise the OM of *S*. Tm. Indeed, uptake of the hydrophobic dye N-phenyl-1-naphthylamine (NPN) was enhanced >5-fold when *S*. Tm was grown in LPM (Figure S2B), a result consistent with increased OM permeability^29^. As predicted, we found that metergoline synergized with several OM-perturbing agents, including EDTA, polymyxin B, and polymyxin B nonapeptide, and was also potentiated in an efflux-deficient strain (Figure 4B and Figure S2C). Intriguingly, the ability of polymyxin B to potentiate metergoline against WT or ∆*tolC S*. Tm was lost in LPM. Low pH, Mg^2+^ (both of which are recapitulated in LPM) and cAMPs are environmental signals for the PhoPQ and PmrAB two-component systems, which regulate lipid A modifications to reduce the affinity of LPS for cAMPs^30^. Therefore, the inefficacy of polymyxin B (≥64-fold increase in MIC) in LPM is unsurprising, with respect to both metergoline potentiation and direct antibacterial activity against *S*. Tm. However, deletion of *phoP* not only restores the synergy between polymyxin B and metergoline in LPM, but also enhances the synergy in MHB (Figure S2C). Additionally, PhoP did not have a major impact on LPM-dependent permeabilization of the OM as determined by NPN uptake (Figure S2B).

We hypothesized that metergoline may be more active against Gram-positive bacteria, which do not possess an OM barrier. As expected, metergoline was >8-fold more potent against methicillin-resistant *Staphylococcus aureus* (MRSA) than *S*. Tm in MHB (Figure S2D), and approximately equipotent against both bacteria in primary macrophages (Figure 3A and Figure S2E). Surprisingly, metergoline still synergized with polymyxin B against MRSA. In addition to OM disruption, polymyxin B can disrupt the cytoplasmic membrane of both Gram-negative and Gram-positive bacteria, and the latter activity is enhanced by the inner membrane activity of bicarbonate^31^. Indeed, bicarbonate increased the potency of metergoline against MRSA and dramatically enhanced the synergy with polymyxin B in both MRSA and *S*. Tm (Figure 4B and Figure S2D).

The above data strongly suggested that the activity of metergoline against intracellular *S*. Tm is in large part due to pH-dependent permeabilization of the OM. We therefore used atomic force microscopy (AFM) to examine the surface topology of intracellular *S*. Tm. We quantified surface roughness of *S*. Tm recently isolated from primary bone marrow-derived macrophages (see Materials and Methods) and compared this to *S*. Tm grown in LPM or MHB (Supplementary Dataset S6). We observed almost a 50% increase in average surface roughness for intracellular *S*. Tm relative to those grown in MHB, and close to a 250% increase in surface roughness for bacteria grown in LPM (Figure 4C and Figure S3). Remarkably, *S*. Tm isolated from macrophages treated with bafilomycin displayed a largely uniform surface topology comparable to cells grown in MHB. This provides direct evidence that the acidic intracellular environment alters the OM of *S*. Tm and offers a compelling explanation for the increased potency of metergoline against macrophage-internalized bacteria compared to cells cultured in standard lab media. When considered with our chemical-genetic interaction data demonstrating the importance of cell envelope homeostasis in macrophages (Figure 1), and the potentiation of conventionally Gram-positive targeting antibiotics we observed in our chemical screens (Supplementary Dataset S3), these results suggest that membrane perturbation of intracellular *S*. Tm potentiates molecules that may otherwise be incapable of traversing the Gram-negative OM.

### Metergoline disrupts the inner membrane and reduces cellular energy pools

As noted above, both active efflux and the OM permeability barrier contribute to the poor activity of metergoline against *S*. Tm grown in MHB. Thus, disruption of OM integrity is required for metergoline activity. Sensitivity to metergoline can be induced *in vitro* through genetic disruption of efflux (i.e. a ∆*tolC* strain). Indeed, we discovered that metergoline is bacteriolytic against a ∆t*olC* strain of *S*. Tm (Figure 4D). In this strain, we were unable to isolate spontaneous resistance mutants after several attempts plating at 4xMIC (observed frequency of resistance <4.8 x 10^−10^). Recently, it has been proposed that antibiotic-induced bacterial cell death can be partially attributed to an imbalance in proton homeostasis across the inner membrane^32,33^. Moreover, previous work suggested that the antifungal activity of metergoline against *Candida krusei* was due to depolarization of the mitochondrial membrane potential and induction of reactive oxygen species^34^. Lastly, inner-membrane active bicarbonate^31^ increased the potency of metergoline against both OM-perturbed *S*. Tm and MRSA (Figure 4B and Figure S2C). Together, these observations led us to hypothesize that metergoline influences membrane potential in *S*. Tm. In Gram-negative bacteria, the inner cytoplasmic membrane maintains the proton motive force (PMF) at a constant value to generate energy that is necessary for ATP synthesis by the F1-F0 ATPase^35^. The PMF is composed of electrical potential (Δψ) and a transmembrane proton gradient (ΔpH), and perturbations to either component result in precise compensatory increases to the other^36^. This process may be targeted by membrane potential-uncoupling antibiotics, wherein the dissipation of either Δψ or ΔpH results in a collapse of the PMF^31,37,38^. Remarkably, we discovered that metergoline caused a rapid release of 3,3’-dipropylthiadicarbocyanine iodide (DiSC_3_(5)) (Figure 4E), a fluorescent probe that accumulates in the inner membrane in a Δψ-dependent manner^39^. Disruption of Δψ results in a release of the DiSC_3_(5) dye and increased fluorescence; thus, our results suggested that metergoline treatment impacts electrical potential in *S*. Tm.

The impact of metergoline on the PMF of the inner membrane suggested that it might modulate cellular ATP levels. Indeed, we found that cellular ATP levels were reduced ~10-fold after a 30 min exposure to 128 μg/mL metergoline (Figure 4F). By comparison, the well-known protonophore carbonyl cyanide m-chlorophenyl hydrazone (CCCP) at a concentration of 32 μg/mL led to a <2-fold in ATP levels in the same time frame (Figure S4A). CCCP selectively targets ΔpH^40^, and we observed striking synergy between metergoline and CCCP, supporting an inner membrane-specific role for metergoline (Figure S4B). We observed a similar effect of metergoline on DiSC_3_(5) release in MRSA (Figure S4C), as well as synergy between three distinct PMF-dissipating molecules with metergoline (Figure S4D).

### Chemical structure-activity-relationship identifies two metergoline analogues with antibacterial activity

Having examined the bacterial determinants for metergoline efficacy, we wanted to explore the chemical properties of the metergoline scaffold required for activity. We therefore tested sixteen structurally related analogs for the ability to reduce growth of wildtype and ∆*tolC S*. Tm in MHB, LPM, and macrophages, as well as MRSA grown in MHB (Table S2). Four of the analogs evaluated herein were FDA-approved therapeutics that contain the core ergot alkaloid scaffold but diverge structurally from metergoline in multiple regions (Figure 5). These compounds: nicergoline (**2**), pergolide (**3**), cabergoline (**4**), and methysergide (**5**), are neuroactive drugs that interact with dopamine and/or serotonin receptors, similar to metergoline (**1**). The remaining twelve analogs were synthesized from metergoline by systematic replacement of the benzyl carbonate moiety with other groups (Figure 5, compounds **6-17**).

**Figure 5.**
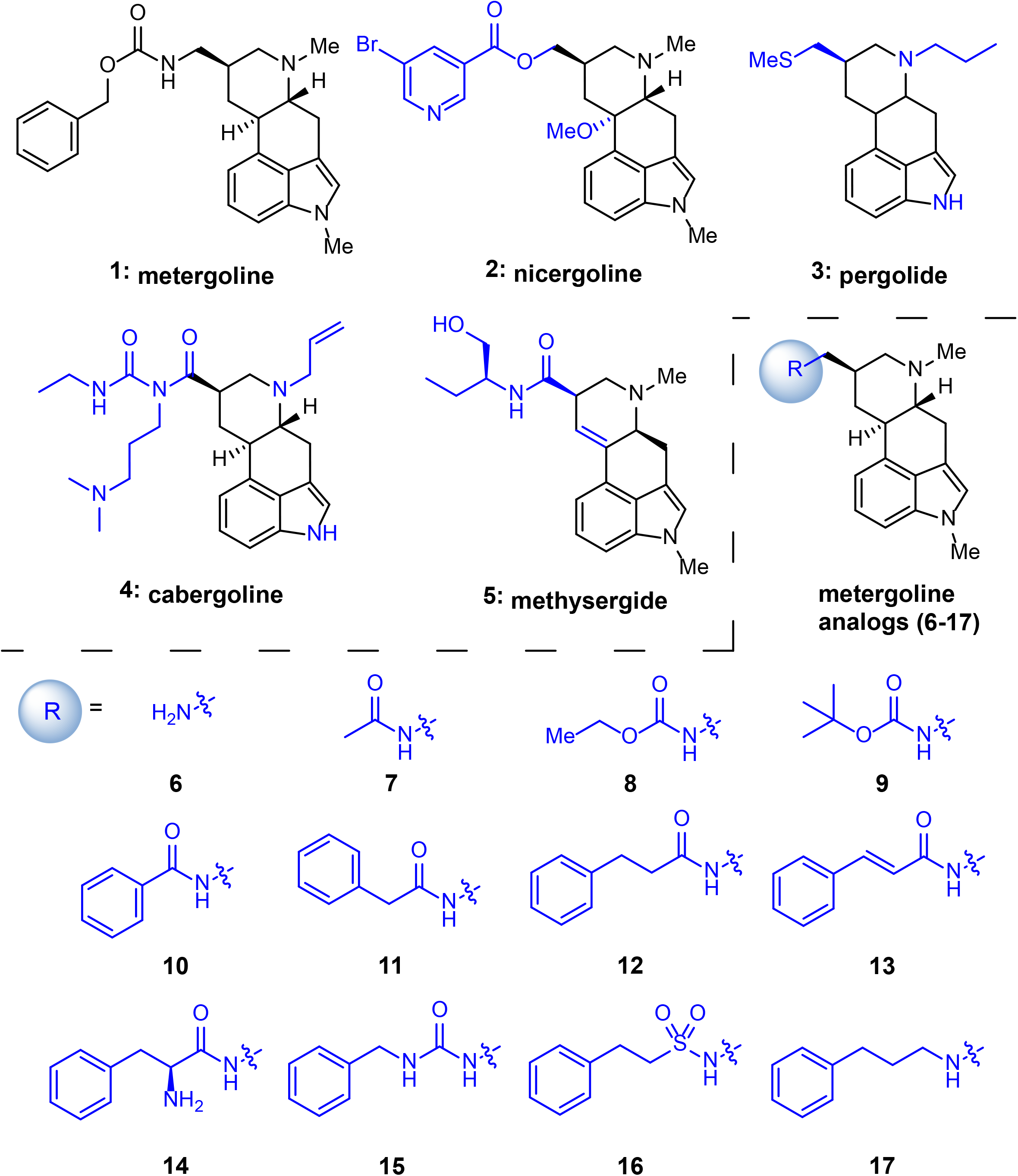
Chemical structures of metergoline and analogs. Structural features that are variations from metergoline are indicated with blue coloring.

Of the four ergot-based drugs, nicergoline (2) showed modest antimicrobial efficacy while the remaining compounds (**3-5**) were inactive in our assays (Table S2). Given that the activity of metergoline appears to be enhanced in the low pH environments of LPM and macrophages, we considered the possibility that metergoline may be acting as a pro-drug by means of hydrolysis of the acid-labile carbamate moiety to release its corresponding amine **6**. Evaluation of compound **6**, however, showed no *in vitro* efficacy at 128 ug/mL in all but one assay (in MHB supplemented with EDTA) and inferior intra-macrophage growth inhibition relative to metergoline (0.39 growth vs. control). The remaining synthetic analogs consisted of: acetamide **7**, alkyl carboamates **8** and **9**, amide derivatives **10-14**, benzyl urea derivative **15**, sulfonamide **16**, and amine **17** (Figure 5). Most of these analogs were inactive in our assays but compounds **13** and **16** demonstrated antimicrobial activity comparable to metergoline. *In vitro*, the α,β-unsaturated amide **13** was twice as potent as metergoline while the phenethyl sulfonamide **16** had approximately half of the potency of metergoline. Both **13** and **16** also inhibited intra-macrophage growth of *S*. Tm with similar potency to metergoline. These compounds were more toxic to macrophages (18.07 and 19.86 % toxicity) relative to metergoline (2.9 % toxicity) as determined by lactate dehydrogenase release (Table S2).

### Metergoline reduces bacterial load in a murine model of *S*. Tm systemic infection

Given its potency *in vitro*, we tested the efficacy of metergoline against a systemic murine infection with *S*. Tm. Mice intraperitoneally infected with *S*. Tm develop symptoms consistent with typhoid fever and typically succumb within 72 hours. Metergoline is a naturally occurring alkaloid compound derived from ergot fungus, and, to our knowledge, its *in vivo* efficacy has been explored solely in experiments to characterize its anxiolytic effects in mice as a 5HT (serotonin) antagonist^41-43^. We found that metergoline (5 mg/kg) significantly reduced bacterial load in the spleen, liver, cecum, and colon (Figure 6A) and significantly extended animal survival time (Figure 6B). As expected, one of the amide derivatives (**12**) which we found to have poor *in vitro* activity did not reduce bacterial load nor prolong survival. These results demonstrate that metergoline has antibacterial activity against *S*. Tm *in vivo*.

**Figure 6.**
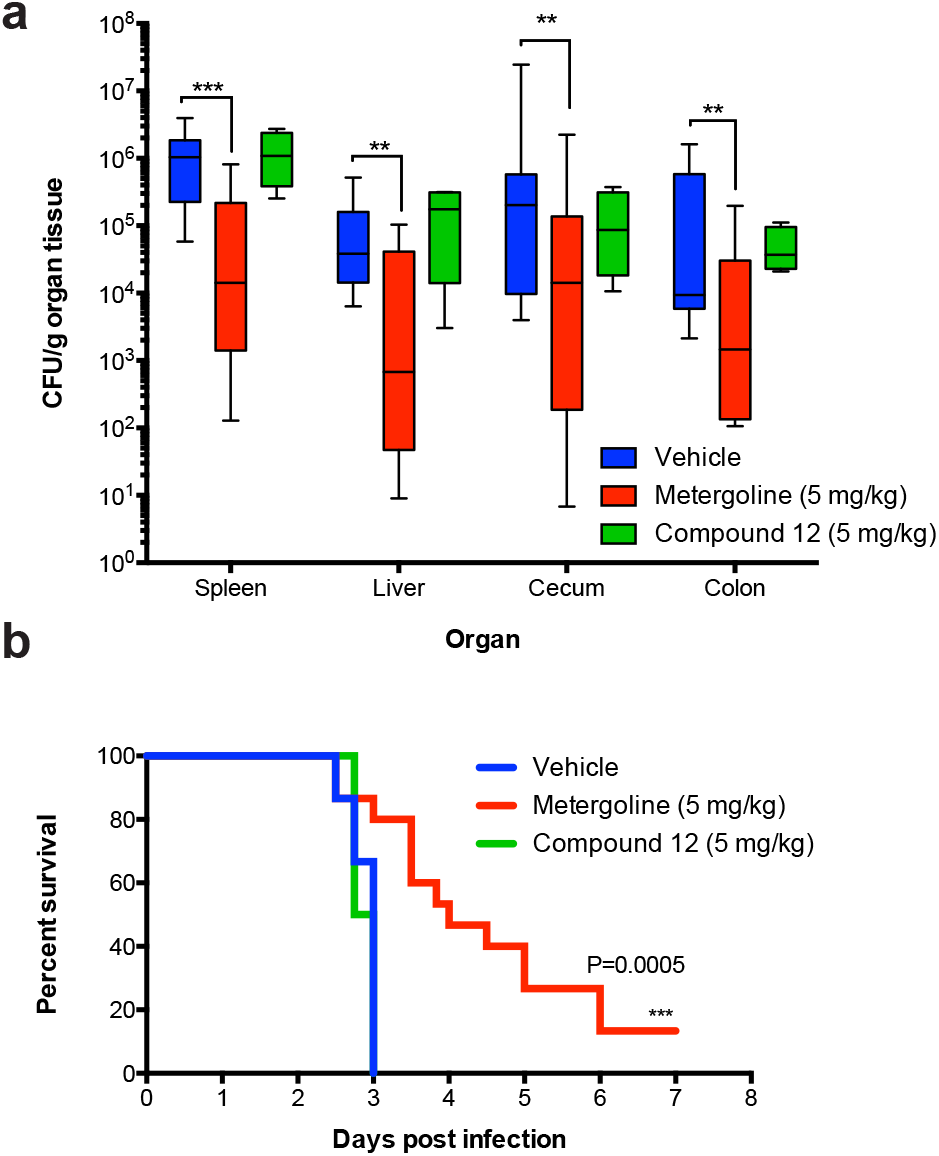
*In vivo* efficacy of metergoline in a murine model of systemic *S*. Tm infection. C51BL/6 mice were infected intraperitoneally with ~10^5^ CFU *S*. Tm. (A) Groups of mice were treated twice daily with metergoline (5 mg/kg, red bars), compound 12 (5 mg/kg, green bars), or DMSO (5% in DMEM, blue bars) by i.p. injection. Mice were euthanized at experimental endpoint (60 hours post infection). Bacterial load in the spleen, liver, cecum, and colon was determined by selective plating on streptomycin. Data shown are the means of three separate experiments (n=5 per group). Box plot whiskers show the minimum to maximum values per group, lines in box plots show the median of each group. Groups were analyzed with a two-way ANOVA and corrected for multiple comparisons with a Holm-Sidak test. *P<0.05, **P<0.01, ***P<0.001, ****P<0.0001. (B) For survival experiments, groups of mice were treated twice daily with metergoline (5 mg/kg, red bars), compound 12 (5 mg/kg, green bars), or DMSO (5% in DMEM, blue bars)) by i.p. injection, and were euthanized at clinical endpoint. Survival curves shown are from three separate experiments (n=5 per group). Groups were analyzed with a Gehan-Breslow-Wilcoxon test for survival curve differences. *P<0.05, **P<0.01, ***P<0.001, ****P<0.0001.

## Discussion

Intracellular pathogens often occupy niches that are difficult to recapitulate *in vitro;* thus, we and others have developed fluorescence, luminescence, and colorimetric-based screening assays with readouts that approximate bacterial viability over traditional cfu counting methods^44-48^. Here we describe a macrophage-based chemical screen specific for intracellular *S*. Tm, leading to the discovery of metergoline as an antibacterial with *in vivo* efficacy, and the unexpected identification of outer membrane disruption in the intracellular environment. Further, we show that metergoline disrupts the proton motive force across the *S*. Tm cytoplasmic membrane, diminishes cellular ATP levels, reduces bacterial load in a murine model of systemic *S*. Tm infection, and significantly prolongs survival of infected animals. This work highlights the ability for intracellular chemical screens to accurately predict *in vivo* efficacy, and their potential to yield better leads against intracellular bacteria. Such screens offer the opportunity to directly assess host cell cytotoxicity to select against off-target effects, and better select molecules with favorable pharmacological properties within host cells, which typically represent considerable challenges in the treatment of intracellular infections.

Our chemical screen was selective for molecules with antimicrobial efficacy conditional to the intracellular environment. We found that metergoline was inactive in standard growth media at neutral pH and that the acidification-blocking activity of bafilomycin abrogated its activity, suggesting that low pH of LPM and/or the SCV sensitizes *S*. Tm to othwerwise inactive compounds. Because we were able to recapitulate the antimicrobial activity of metergoline in non-acidic environments by inducing OM perturbation with membrane-permeabilizing agents (EDTA, PMB) or in a ∆*tolC* mutant background, we sought to characterize OM integrity in *S*. Tm grown under acidic conditions. Low pH in the intracellular environment is a critical signal for multiple two-component regulatory systems in *S*. Tm (PhoPQ, PmrAB, SsrAB)^49-51^, such that its detection acts as a cue for virulence gene expression. Given the importance of low pH as a regulatory input governing *S*. Tm pathogenesis within cells, antibiotics selective for intracellular *S*. Tm must account for the genetic and physiological profiles of *S*. Tm grown in acidic conditions. Interestingly, we found that acidified *S*. Tm undergoes a loss of normal OM surface structure and experiences OM disruption, as quantified by NPN uptake. These data strongly suggest that acidic pH in the intracellular environment perturbs the OM. Our observation that conventionally Gram-positive antibiotics are potentiated in LPM and macrophages provides further evidence for increased OM permeability in acidic environments. Taken together, these findings have important implications for antibiotic drug discovery. Gram-negative bacteria are susceptible to antibiotics capable of directly diffusing through the OM lipid layer (e.g. polymyxins and aminoglycosides) and small hydrophilic antibiotics that traverse the OM through porin channels ^52,53^. Overcoming the Gram-negative OM permeability barrier is currently one of the greatest challenges in drug-discovery programs directed at these difficult to treat infections. However, our work indicates that natural components of the innate immune system may sensitize intracellular Gram-negative bacteria to otherwise poorly penetrating antibacterial compounds *in vivo*.

Overall, we propose a model for metergoline’s antibacterial activity *in vivo* wherein this compound is capable of accumulating in the SCV, then targets the cytoplasmic membrane due to pH-dependent permeabilization of the *S*. Tm OM. It is tempting to speculate that the low pH in the SCV enhances metergoline’s activity by altering the relative contributions of membrane potential (Δψ) and pH gradient (∆pH) to the proton motive force, however, further work is required to investigate this. Screening for antimicrobials selective for bacteria within macrophages is able to select for those that target conditionally essential genes, due to the unique conditions of the intracellular environment. Indeed, genes that are essential for growth under conditions that more closely resemble those during infection is a promising avenue in modern antibiotic research that has garnered increased attention in recent years ^54^. For complex *S*. Tm infections that include several bacterial subpopulations with highly varied gene expression and virulence phenotypes, the identification of novel antimicrobials with unique targets is critical. We conclude that intracellular-targeting chemical screening platforms have the potential to identify small molecules with relatively underexplored targets, a goal of paramount importance in the current antibiotic resistance era.

## Acknowledgments

We are grateful to Susan McCusker for excellent technical assistance in high-throughput compound screens, and Maya Farha for useful discussions throughout this work. M.J.E was supported by a post-doctoral Fellowship from the Canadian Institutes of Health Research (CIHR); C.N.T. was supported by a Canada Graduate Scholarship from the Natural Sciences and Engineering Research Council and an Ontario Graduate Scholarship. This work was supported by operating grants from CIHR to B.K.C (388221 and 316614) and E.D.B (FRN-143215)), the Canada Research Chairs program (to B.K.C. and E.D.B), the Ontario Research Fund (to B.K.C., and E.D.B.), and a donation from the Boris Family Fund for Health Research Excellence (RE01-048 to B.K.C. and E.D.B.). J.W.J. and J.M. were supported by a startup grant from the McMaster Faculty of Health Sciences Dean’s Fund. M.M., S.P. and H.A.P. were supported in part by NIH grant R01AI075093, and NIAID Contract No. HHSN212200900040C.

## Author Contributions

M.J.E., C.N.T., B.K.C., and E.D.B conceived and designed the research. J.W.J and J.M designed metergoline analogues. M.J.E. and C.N.T. performed all experiments and analyzed data with the following exceptions: J.W.J. synthesized and analyzed metergoline analogues, S.F. performed atomic force microscopy, W.E. assisted with mechanistic studies. S.P., H.A-P., and M.M. constructed and supplied the *Salmonella* deletion library. M.J.E., C.N.T., B.K.C., and E.D.B wrote the paper. All authors commented on the manuscript.

## Data Availability Statement

Source data for Figures 1, 2, and 4 are provided as Supplemental Datasets with the paper.

## MATERIALS AND METHODS

### Ethics statement

All animal experiments were performed according to the Canadian Council on Animal Care guidelines using protocols approved by the Animal Review Ethics Board at McMaster University under Animal Use Protocol #11-03-10.

### Chemicals

Screening stocks (5mM) of the Pharmakon-1600 (MicroSource) compound library were stored at −20°C in DMSO. With the following exceptions, all chemicals were purchased from Sigma-Aldrich: Ciclopirox (Santa Cruz Biotechnology), Pergolide (Toronto Research Chemicals), Cabergoline (Toronto Research Chemicals), Methysergide (Toronto Research Chemicals). Compounds were routinely dissolved in DMSO at a concentration of 12.8 or 25.6 mg/mL and stored at −20°C. Synthetic analogs of metergoline are described in Supplementary Information.

### Bacterial strains and culture conditions

All experiments with *Salmonella enterica* subsp. *enterica* ser. Typhimurium (*S*. Tm) were performed with strain SL1344 or derivatives with the exception of the *S*. Tm single-gene deletion collection ^19^, which was constructed in the related *S*. Tm str. 14028s background. For consistency, in the experiments presented in Figure 1 and Figures S1 and S3, the wildtype *S*. Tm 14028s strain was used for comparison. For compound screening and secondary assays, *S*. Tm SL1344 was transformed with pGEN-*lux* ^55^, and SL1344 ∆*tolC* was generated by standard methods ^56^. Briefly, the chloramphenicol resistance cassette from pKD3 was amplified by PCR with primers 5’-CAATTATTTTTACAAATTGATCAGCGCTAAATACTGCTTCACAACAAGGAGTAGGCTGGAGCTGC-3’and 5’-AGACCTACAAGGGCACAGGTCTGATAAGCGCAGCGCCAGCGAATAACTTACATATGAATATCCTCCTTAGTTCC-3’ and the resulting amplicon was transformed into electro-competent *S*. Tm SL1344 cells harboring the pSim6 plasmid as previously described ^57,58^. Successful recombinants were confirmed by colony PCR using primers 5’CGACCATCTCCAGCAGCCAC-3’ and 5’GAAAAGGCGAGAATGCGGCG-3’. Experiments with *Staphylococcus aureus* used a Canadian isolate (CMRSA10) of community-acquired methicillin-resistant *S. aureus* USA300 ^59^.

Overnight cultures of bacteria were inoculated with a single colony and routinely grown in LB media (10 g/L NaCl, 10 g/L Tryptone, 5 g/L yeast extract) supplemented with antibiotics as appropriate (streptomycin, 150 μg/mL; chloramphenicol, 25 μg/mL; kanamycin, 50 μg/mL, ampicillin, 100 μg/mL). Where indicated, bacteria were subcultured 1:100 and grown to mid-log phase in cation-adjusted MHB (BBL™ Mueller Hinton II Broth – Cation Adjusted), LPM (5 mM KCl, 7.5 mM (NH_4_)_2_SO_4_, 0.5 mM K_2_SO_4_, 10 mM Glucose, 49 μM MgCl_2_, 337 μM PO_4_^3-^, 0.05% casamino acids, 80 mM MES, pH 5.8), or MOPS glucose minimal medium (Teknova) supplemented with 40 μg/mL histidine when growing *S*. Tm SL1344. Bacteria were grown at 37°C.

### Genetic Screening

The *Salmonella* single-gene deletion (SGD) library ^19^ was pinned from frozen DMSO stocks at 384-colony density onto LB agar medium containing 50 μg/mL kanamycin using a Singer RoToR HDA (Singer Instruments) and grown for 18 h at 37°C. The SGD was then grown overnight in 384-well clear flat-bottom plates (Corning) in either MOPS glucose (Figure 1A) or LB (Figure 4A) supplemented with 50 μg/mL kanamycin. The Singer RoToR HDA was then used to inoculate assay plates (containing 50 μL/well LPM without casamino acids, MHB, or MHB with 100 μg/mL metergoline) with a starting inoculum of approximately 1.7 x 10^5^ CFU/well. Optical density at 600 nm (OD_600_) was measured with a Tecan M1000 Infinite Pro plate reader at the time of inoculation (T_0_) and after 16 hrs (T_16_) incubation at 37°C with shaking at 220 rpm. Growth was calculated by subtracting the pre-reads (T_16_ – T_0_) and normalizing for plate and well effects as previously described ^60^. The interaction score was calculated by dividing normalized growth in the presence of metergoline by the MHB control. Experiments were performed in duplicate or triplicate as noted.

### Replication of SGD mutants in macrophages

Strains from the *Salmonella* single-gene deletion library were selected based on prioritization from genetic screening in LPM (growth < 3.5σ from mean of screening data), along with SPI-1, SPI-2, virulence, motility, and regulatory genes, and WT to use as controls. Strains of interest were grown overnight in 96-well clear flat-bottom plates (Corning) in LB in duplicate.

RAW264.1 macrophages were seeded into 96-well plates in DMEM + 10% FBS at ~10^5^ cells per well and left to adhere for 20-24 hours, incubated at 31°C with 5% CO_2_. Overnight cultures of bacteria were diluted to obtain an MOI of ~50:1, then opsonized for 30 min in 20% human serum in PBS at 31°C. Bacteria were added to each well, and plates were spun down at 500xg for 2 min, then incubated for 30 min at 31°C with 5% CO_2_. Media was aspirated and replaced with fresh DMEM containing 100 μg/mL gentamicin to kill extracellular bacteria for 30 min at 31°C with 5% CO_2_. For half the plates, RAW264.7 cells were washed once with PBS, then scraped from the wells and lysed in PBS containing 1% (v/v) Triton-X100, 0.1% (w/v)

SDS. Bacterial colony-forming units (CFUs) from each well were enumerated by serially diluting in PBS and plating on LB plates (T_0_ counts). For the other half of the plates, RAW cells were washed once with PBS, then fresh DMEM was added and cells were incubated at 31°C with 5% CO_2_. After 7 h, RAW cells were washed once with PBS, then scraped from the wells and lysed in PBS containing 1% (v/v) Triton-X100, 0.1% (w/v) SDS. Bacterial colony-forming units (CFUs) from each well were enumerated by serially diluting in PBS and plating on LB plates (T_7_ counts). Colony-forming units (cfu) were averaged (two technical replicates per assay plate) and a 7h/0h cfu ratio was calculated to represent fold replication over the course of the experiment.

### High-throughput compound screening

For chemical screening in LPM, an overnight culture of *S*. Tm SL1344 was subcultured 1:100 in LB, grown to an OD_600_ of 0.5, then diluted 40-fold into LPM and grown to an OD_600_ of 0.3 before a final 1:150 dilution into LPM. Bacterial culture (50 μL) was dispensed into 96-well black, clear flat-bottom (Corning) plates and then 50 μL of each compound (diluted to 20 μM in LPM) was added for a final concentration of 10 μM compound and ~2×10^4^ CFU/well. The OD_600_ was read immediately after compound addition (T_0_) and then 16 hrs later (T_16_). Plate and well effects were normalized as previously described ^60^ and compounds reducing growth more than 3σ below the mean were considered primary screen actives. Screening was performed in triplicate.

For the screen against intramacrophage *S*. Tm, RAW264.1 macrophages were seeded 16 hrs prior to infection at ~2 x10^5^ cells per well in 96-well black, clear flat-bottom plates (Corning) in DMEM + 10% FBS and were incubated at 31°C with 5% CO_2_. *S*. Tm str. SL1344 transformed with pGEN-lux was grown overnight in LB with 100 μg/mL ampicillin, diluted to obtain a multiplicity of infection (MOI) of 100:1, then opsonized for 30 min in 20% human serum (Innovative Research) in PBS at 31°C. An equal volume of bacteria (100 μL) was added to macrophages and plates were centrifuged at 500 x g for 2 min followed by a 30 min incubation at 31°C with 5% CO_2_. Media was then aspirated and replaced with fresh DMEM containing 100 μg/mL gentamicin to eliminate extracellular bacteria, and plates were incubated for 30 mins at 31°C with 5% CO_2_. Infected cells were washed with PBS prior to addition of 100 μL of DMEM containing compound at 10 μM. Luminescence was read immediately after compound addition (T_o_) and plates were then incubated for 3 hr at 31°C with 5% CO_2_. Luminescence was measured a second time after media was replaced with fresh DMEM + 10% FBS (T_3_) and plates were incubated for a further 3hr before luminescence was measured a final time (T_Final_).

### Secondary screening of compounds in macrophages

130 priority compounds were selected based on prioritization from chemical screening in LPM and macrophages. Infection assays in RAW264.7 macrophages were performed as described above, with the following modifications: macrophages were seeded at 10^5^ cells per well, macrophages were pretreated with 100 ng/mL LPS from *Salmonella enterica* serovar Minnesota R595 (Millipore), and an MOI of 50:1 was used. Compounds were serially diluted two-fold starting at 5 mM to achieve a final concentration of 50 μM with 1% DMSO in DMEM. Luminescence was read immediately after compound addition (t0), and plates were incubated for 20 h at 37°C with 5% CO_2_ before luminescence was read again (t20). Luminescence readings were averaged (two technical replicates per assay plate) and a 20h/0h luminescence ratio was calculated to represent fold replication over the course of the experiment. Minimal inhibitory concentrations (MICs) were estimated based on ability to reduce fold-change in luminescence to 10 or lower.

### Cytotoxicity assays

RAW264.7 macrophages were seeded into 96-well plates in DMEM + 10% FBS and 100 ng/mL *Salmonella enterica* serovar Minnesota R595 (Millipore) and left to adhere for 20-24 hours, incubated at 37°C with 5% CO_2_. Compounds were premixed into DMEM at a final concentration of 50 μM with 1% DMSO, then added to wells. After 2 hours of compound treatment, the culture supernatant was collected for analysis of lactate dehydrogenase release. Cytotoxicity was quantified colorimetrically with the Pierce LDH cytotoxicity kit. Lysis control wells were treated with 10X lysis buffer for 1 hour. Percent cytotoxicity was calculated with the formula: (experimental LDH – spontaneous LDH)/(maximum LDH – spontaneous LDH) x 100, where spontaneous LDH is the amount of LDH activity (read at OD_490_, background subtracted OD_680_) in the supernatant of untreated cells and maximum LDH is the amount of LDH activity in the supernatant of lysis control wells.

### Bone marrow-derived macrophage assays

Bone marrow-derived macrophages (BMMs) were collected from the femur and tibia of 6-10 week old female C57BL/6 mice (Charles River Laboratories) and differentiated in RPMI (Gibco) + 10% FBS + 10% L-sup (L929 fibroblast conditioned medium) + 100 U penicillin-streptomycin for 7 days at 37°C and 5% CO_2_. Differentiated BMMs were seeded 20-24 h prior to infection in 96-well plates at 10^5 cells per well in RPMI + 10% FBS + 100 ng/mL *Salmonella enterica* serovar Minnesota R595 (Millipore) and incubated at 37C with 5% CO_2_. Infection assays were performed as described for RAW264.7 macrophages, with the following modifications: bacteria were not opsonized prior to infection, and an MOI of 50:1 was used. Compounds were added to wells as described previously (see Secondary screening of compounds in macrophages). In experiments where bafilomycin treatment was used, cells were pretreated with 10 μM bafilomycin A-1 or 1% (v/v) DMSO as a vehicle control, for 60 min prior to infection.

### Chequerboard analyses and compound potency analysis

A single colony of freshly streaked bacteria was used to inoculate LB with appropriate antibiotics (150 μg/mL streptomycin for *S*. Tm str. SL1344 and derivatives, 25 μg/mL chloramphenicol for SL1344 ∆*tolC* or ∆*phoP*). Overnight cultures were diluted 100-fold into antibiotic-free MHB or LPM, as appropriate, and grown to mid-log phase (OD_600_ = 0.4-0.8). Subcultures were then diluted to an OD_600_ = 0.0001 (~1×10^5^ cfu/mL) in assay media and 150 μL of this dilution was added to each well of the 96-well assay plate. For chequerboard experiments, an 8×12 matrix of two compounds was created with two-fold serial dilutions of each compound. Metergoline, CCCP, nigericin, and valinomycin were dissolved in DMSO, and polymyxin B, polymyxin B nonapeptide, and EDTA were dissolved in water. After addition of bacteria, plates were incubated at 31°C with shaking for 16 hours, at which time the OD_600_ was measured. The minimum inhibitory concentrations (MIC) for various compounds were determined using 11 two-fold dilutions and growth was measured after 16 hours. The MIC was the concentration that inhibited growth >95% when compared to the solvent control. In all experiments, DMSO was present at a final concentration <2% (routinely 1%).

### NPN Uptake Assay

Uptake of the lipophilic dye N-phenyl-1-naphthylamine (NPN) was measured essentially as previously described ^29^. Briefly, overnight cultures of *S*. Tm were diluted 50-fold into LB and grown to mid-log (OD_600_ = 0.5), then diluted again 100-fold into LPM or MHB and grown to late-exponential phase (OD_600_ ~1.2). Cells were harvested by centrifugation, washed in 5 mM HEPES, pH 1.2, then resuspended to a final OD_600_ = 1.0. The cell suspension (50 μL), 40 μM NPN (50 μL), and 5 mM HEPES, pH 7.2 (100 μL) were mixed in black clear flat-bottom 96-well plates immediately before measuring fluorescence (excitation, 340 nm; emission 415 nm) in a Tecan M1000 Infinite Pro plate reader. NPN uptake is reported relative to maximum fluorescence (obtained by presence of 50 μg/mL polymyxin B) after background (cells and buffer without NPN) subtraction.

### Atomic force microscopy (AFM)

BMMs were differentiated as described above, and then seeded 16 hours prior to infection in 6-well plates at 5 x 10^6 cells per well in RPMI + 10% FBS + 100 ng/mL *Salmonella enterica* serovar Minnesota R595 (Millipore) and incubated at 31°C with 5% CO_2_. Cells were pretreated with 10 μM bafilomycin A-1 or 1% (v/v) DMSO as a vehicle control, for 60 minutes prior to infection. Bacteria were added at an MOI of 100:1 and were allowed to infect for 30 minutes, after which media was aspirated and replaced with fresh RPMI containing 100 μg/mL gentamicin (to kill extracellular bacteria) for 30 minutes at 31°C with 5% CO_2_. Media was then aspirated and replaced with fresh RPMI, and macrophages were scraped into the media.

20 μL of suspended macrophages was then transferred to a hydrophilic polycarbonate 0.2 μm Millipore Isopore GTTP filter (Merck Millipore), with a Kimwipe (Kimberly-Clark Professional) underneath to remove liquid without vacuum. Filters were attached to a glass slide with an adhesive and examined using AFM. Macrophages lysed immediately upon removal of medium, with bacterial cells remaining intact for surface scanning. For bacteria grown in various growth media, cells were grown to mid-log phase (OD_600_ ~ 0.5), then 20 μL of the suspension was placed on the same filter as above. The liquid was removed with a Kimwipe as previously described, then 20 μL of 10 mM MES pH 5.5 was overlaid, and liquid removed. Filters were mounted on a glass slide and examined with AFM.

A Bruker BioScope Catalyst AFM, with a Nanoscope V controller, was used to scan bacterial surfaces. For each sample, a 0.65 μm thick Si3N4 triangular cantilever was used (Scan Asyst AIR, Bruker), with a symmetric tip and spring constant of ~0.4 N·m^−1^. All AFM was done at 25°C (ambient room temperature), with a scan rate of 0.5 Hz and 256 samples per line resolution. Scanning was done in PeakForce quantitative nanomechanical mapping mode. All downstream image processing was done using NanoScope software (Bruker). For scans of whole cells, scans were fit to a plane to normalize the Z-Height. For scans of bacterial surface topology, images were flattened using a second order transformation to account for subtle cell curvature, and surface topography was calculated from cross sections of these image scans.

### DiSC_3_(5) assay

Subcultures of WT *S*. Tm or MRSA were grown to late-exponential phase (OD_600_ ~ 1) in MHB (MRSA) or MHB with 10 mM EDTA (*S*. Tm). Gram-negative outer membrane disruption is required for the highly lipophilic 3’3-dipropylthiadicarbocyanine iodide (DiSC_3_(5)) to access the cytoplasmic membrane ^61^. Cells were harvested by centrifugation, washed twice in buffer (5 mM HEPES, pH 7.2, 20 mM glucose), and then resuspended in buffer to a final OD_600_=0.08 5 with 1 μM DiSC_3_(5). After a 20 min incubation at 37°C, 150 μL of DiSC_3_(5) loaded cells was added to two-fold dilutions of metergoline in 96-well black clear-bottom plates (Corning) and fluorescence (excitation = 620 nm, emission = 685 nm) was read 1 min later using a Tecan M1000 Infinite Pro plate reader. The fluorescence of metergoline diluted in buffer was negligible (<200 a.u.). The fluorescence intensity was stable (<5% fluctuation) for at least 15 min when the plate was shielded from light.

### Measurement of intracellular ATP levels

WT *S*. Tm was grown in MHB with 1 mM EDTA to early-log phase (OD_600_ = 0.2) and then grown in the presence of metergoline for 30 minutes in clear flat-bottom 96-well plates. The OD_600_ was determined immediately before ATP levels were measured using a BacTiter-Glo™ Microbial Cell Viability Assay (Promega), according to manufacturer instructions, in a white 96-well plate using an EnVision plate reader (PerkinElmer). Relative ATP levels were calculated by dividing relative light units (RLU) by the OD_600_ (RLU/OD).

### Mouse infections

Before infection, mice were relocated at random from a housing cage to treatment or control cages. Six- to ten-week-old female C57BL/6 mice (Charles River Laboratories) were infected intraperitoneally with ~10^5^ cfu *S*. Typhimurium SL1344 in 0.1 M Hepes (pH 8.0) with 0.9% NaCl. Metergoline was administered at 5 mg/kg via intraperitoneal injection, solubilized in 5% DMSO in DMEM. 5% DMSO in DMEM was given to vehicle control groups of mice. Clinical endpoint was determined using body condition scoring analyzing weight loss, reduced motility, and hunched posture. Experimental endpoint was defined as 60 hours post-infection for cfu comparison experiments. At experimental endpoint, mice were euthanized, and the spleen, liver, cecum, and colon were aseptically collected into ice-cold PBS and homogenized. Bacterial load in each tissue type was enumerated from organ homogenates serially diluted in PBS and plated onto solid LB supplemented with 100 μg/mL streptomycin.

### Statistical analysis

Data were analyzed using R or GraphPad Prism 6.0 software (GraphPad Inc., San Diego, CA), with statistical tests indicated in figure legends. P values of <0.05 were considered significant.

